# Infectivity of three Mayaro Virus (Genus Alphavirus, Family Togaviridae) geographic isolates in human cell lines

**DOI:** 10.1101/2021.09.29.462323

**Authors:** Aum R. Patel, Melissa Dulcey, Nabil Abid, Melanie N. Cash, Jordan Dailey, Marco Salemi, Carla Mavian, Amy Y. Vittor

## Abstract

Mayaro virus (MAYV) is an emergent arthropod-borne virus that causes an acute febrile illness accompanied by arthralgia, similar to chikungunya virus. Increasing urbanization of MAYV outbreaks in the Americas has led to concerns that this virus could further expand its geographic range. Given the potential importance of this pathogen, we sought to fill some critical gaps in knowledge regarding MAYV infectivity and geographic variation. This study describes the cytopathogenicity of MAYV in human dermal fibroblasts, human skeletal muscle satellite cells, human embryonic kidney cells (HEK), peripherally derived human macrophages, and Vero cells. MAYV strain isolated from Bolivia (MAYV-U) infected cells more rapidly compared to MAYV strains isolated in Peru and Brazil (MAYV-P; MAYV-B), with high titers (1×10^8^ pfu/ml) peaking at 37 hours post infection. MAYV-U also caused the most cytopathic effect in a time dependent manner. Furthermore, differently from the other two prototypic strains, MAYV-U harbors unique mutations in the E2 protein, D60G and S205F, likely to interact with the host cell receptor, which may explain the observed differences in infectivity. We further demonstrate that pre-treatment of cells with interferon-β inhibited viral replication in a dose-dependent manner. Together, these findings advance our understanding of MAYV infection of human target cells and provide initial data regarding MAYV phenotypic variation according to geography.

**Author Summary:** Arthropod-borne viruses are of great public health concern, causing epidemics worldwide due to climate change, changes in land use, rapid urbanization, and the expanding geographic ranges of suitable vectors. Among these viruses, Mayaro is an emerging virus for which little is currently known. This study aims to answer fundamental questions of Mayaro virus biology using three geographically distinct viral strains to examine variability in infection kinetics and infectivity in susceptible cell types. We found one geographic isolate to have accelerated infection kinetics and increased cell damage because of infection. To better understand what was unique about this isolate, we compared their envelope protein, which is critical for entry into a cell. We found that the isolate with increased replication kinetics possessed mutations at sites that may promote viral entry, which could explain these findings. Together, these findings further our understanding of Mayaro virus biology and provide insight into factors that contribute to Mayaro transmission and infectivity.

## Introduction

Mayaro virus (genus *Alphavirus*, family *Togaviridae*; MAYV) is an emerging arthropod-borne virus transmitted by *Haemagogus* mosquitoes in forest regions of Central and South America (1) and the Caribbean (2,3). Similar to other Old World alphaviruses such as chikungunya virus (CHIKV), MAYV infection leads to fever, maculopapular rash, and persistent arthralgias (4). In most symptomatic cases, alphavirus infections are acute and self-limited. However, reports of lasting arthritis, neurological complications, myocarditis and even death have also been reported (5–8). Despite this, neither MAYV epidemiology nor virology is well characterized (5–8). A recent publication by *Bengue* et al. characterized MAYV infection in human chondrocytes, fibroblasts, and osteoblasts by examining replication kinetics and upregulation of proinflammatory cytokines and arthritis related genes due to infection (9). Their study, along with ours, aims to further our understanding of MAYV pathogenesis and the risk it poses to human health.

MAYV was first isolated in Trinidad and Tobago in 1954 (3). It has been found in several countries following its discovery, including Bolivia (10), French Guiana (11), Peru (12), Venezuela (13), and Brazil, with seropositivity in endemic areas such as Brazil ranging from 5% to as high as 60% (4). An increasing number of MAYV infections have been reported from peri-urban areas signaling a possible adaptation of the virus from a sylvatic to a grassland ecology, indicating a potential expansion of its geographic range (14). This expansion is further supported by the recent detection of MAYV in Haiti and Mexico (2,14,15).

Phylogenetic analysis classifies MAYV into three genotypes: D (dispersed, MAYV-D) and L (limited, MAYV-L) (4), named on the basis on their geographic distribution, and the N (new) genotype recently identified (1). L/D hybrid genotypes have also been described, which emerged from recombination events in the Amazon basin (16). The main D and L genotypes have coexisted for decades in the Pan-Amazonian region and are hypothesized to use sloths, marmosets, and other non-flying mammals to help sustain their independent enzootic transmission cycles (17). Populations infected with each genotype did experience viral flow (migration) events during their coexistence, yet their genomic sequences remain highly similar to one another, suggesting strong selective negative/purifying pressure that would hindering significant divergence over time (17). While phylogeny and evolutionary dynamics of such genotypes have been well characterized, no studies have investigated to our knowledge how the existing, albeit small, genetic diversity may impact pathogenesis or viral fitness in either *in vitro* or *in vivo* model systems.

Our aim is to characterize the infectivity of distinct geographic MAYV isolates in human cell lines and to identify specific features of the envelope polyprotein sequences that may contribute to any differences seen. We predict that these differences, if they exist, may result from variability in the intrinsic cellular antiviral response, viral entry, post entry barriers to productive infection, or any combination of these factors. This study also examines the MAYV envelope protein to determine whether selective pressures have induced diversifying selection in key regions relevant to viral binding and target cell infection. Together, our findings serve as part of the foundation to better understand MAYV pathogenesis, host/vector susceptibility, and geographic expansion, which are critical for characterizing MAYV and the risk it poses.

## Methods

### Cells

Vero E6 cells (ATCC) and HEK293T cells (kindly provided by Dr. David Pascual) were cultured at 37°C with 5% CO2. Both were maintained in Corning Dulbecco’s modified Eagle’s medium (DMEM) with L-Glutamine, 4.5g/L Glucose and Sodium Pyruvate, supplemented with 5% EqualFETAL™(Atlas Biologicals, Fort Collins, CO), 1% Penicillin-Streptomycin (Thermo Fisher Scientific, Waltham, MA) and an additional 1% L-Glutamine (Thermo Fisher Scientific, Waltham, MA).

Normal Adult Human Dermal Fibroblasts and Normal Human Skeletal Muscle Satellite Cells (SkMc) (Lifeline Cell Technologies, USA) were cultured using FibroLife S2 Medium and StemLife SK Medium, respectively.

To obtain macrophages, peripheral blood monocytes (PBMCs) were isolated from the whole blood of healthy donors using an EasySep Direct Human PBMC Isolation Kit (StemCell Technologies) through negative selection. Following isolation, PBMCs were plated in flasks for monocyte adhesion and the M1 Macrophage Differentiation Kit (PromoCell, USA) was used to differentiate macrophages from monocytes.

### Ethics

Ethics approval for PBMC isolation from healthy blood donors was obtained from the University of Florida Institutional Review Board (IRB 201600448). Study participation was voluntary, and written informed consent was obtained from adults (18 years of age and older). Only adult (age of 18 years and older) participants were involved.

### Virus and Propagation

Mayaro virus (1955 Uruma, Bolivia) designated as MAYV-U was purchased from BEI Resources (NR-49914, USA). Mayaro virus (2000 Loreto, Peru) designated as MAYV-P and Mayaro virus (1955 Para, Brazil) designated as MAYV-B were obtained from World Reference Center for Emerging Viruses and Arboviruses (WRCEVA; MAYV IQU3056 and MAYV BeH407 respectively). These viruses were all isolated from humans.

Each vial of virus was resuspended in DPBS without Ca/Mg and propagated in Vero cells in DMEM prepared as previously described. Supernatants from the infected cell culture were subsequently harvested 48, 72, and 96 hours (hrs) from MAYV-U, MAYV-P, and MAYV-B isolates respectively, centrifuged at 1200 xg for 15 minutes to remove any remaining cell debris, and frozen at -80°C. Prior to use, vials containing virus were thawed once at 37°C and never refrozen.

### MAYV Titration and Plaque Assay

Serial dilutions (10-fold) of harvested virus supernatant (200 uL) were added to individual wells in duplicate to Vero cells grown to 100% confluence in 12-well plates. An additional 200 uL of prepared DMEM was added to each well to prevent the cells from drying out. The plates were incubated at 37°C, 5% CO_2_ for 1 hr to allow for viral adsorption before the media was aspirated and replaced with 0.5 ml of 1% methyl-cellulose medium (5% EqualFETAL™, 2% Pen Strep, 2% L-Glut in 2x E-MEM). Cells were incubated for 3 days before the media was removed and stained with 0.25% crystal violet (30% methanol; 70% dH_2_O) for 30 minutes. The crystal violet was then aspirated and the plates were washed with water. Plaques were counted manually and the concentration in plaque-forming units/ml (pfu/ml) was calculated.

### Immunofluorescence Microscopy

Cells were grown on 12mm culturable coverslips coated with Poly-D-Lysine (Neuvitro, Camas, WA). Once cells reached 80% confluence, they were infected at a multiplicity of infection (MOI) of 1 for 24, 48, and 72 hrs for immunofluorescent staining. Infected media was aspirated and the cells were fixed with 4% paraformaldehyde (PFA) for 15 minutes. The PFA was aspirated and the cells gently washed with DPBS w/o Ca/Mg. To permeabilize cell membranes, 0.1% Triton X-100 was added for 15 minutes, and then removed before the cells were washed again as before.. Triton X-100 was removed and the cells were washed again. To block nonspecific antibody binding, 0.5 ml of 3% BSA in DPBS was added for 1 hr. For viral antigen staining, the primary antibody, Anti-Eastern Equine Encephalitis clone 1A4B-6 Mab IgG2b was added to the cells at a dilution of 1:1000 and incubated for 1 hour. This antibody reacts with an E1 epitope shared by all alphaviruses. Next, the cells were washed with DPBS before an hour incubation with a 1:1000 dilution of Goat anti-Mouse IgG (H+L), Alexa Fluor 488 (Invitrogen, Carlsbad, CA).

Following incubation, the secondary antibody was aspirated, and DAPI at a dilution of 1:1000 was added for 15 minutes. To view the cells, coverslips were mounted to a microscope slide using Prolong TM Gold Antifade Mountant (ThermoFisher, Waltham, MA). Microscope slides were imaged using an Olympus IX-81 DSU Spinning Disk Confocal microscope under 20x, using Z-Stack and imaged without gamma. Images were edited by creating projection images from the Z-Stacks, which were then deconvoluted and re-normalized. Exposure conditions were kept consistent within a microscopy session.

### Viral Replication Kinetics and Cytopathic Effect Staining

Cells were grown in T-25 flasks to 100% confluence. One flask was used for each time point and viral isolate. A total of 13 time points at 6-hr intervals from 0-72 hrs (with uninfected controls) were performed in duplicate. Before inoculation at time zero, the growth media was removed and replaced with fresh media. Cells were then infected at a MOI of 1 and incubated at 37°C, 5% CO_2_ for a 1 hour adsorption period. Uninfected cells received fresh media and were incubated for the same amount of time. Following incubation, the media was removed from all flasks, the cells were washed with DPBS w/o Ca/Mg, and then replenished with fresh media. At the corresponding time points, the media was collected by aspiration and placed individually in 15 mL centrifuge tubes. Tubes were then centrifuged at 1200 xg for 15 minutes to remove any remaining cell debris, and cryovials were prepared using the supernatant, which were subsequently stored at -80°C. To determine viral titer, one vial of frozen supernatant was thawed, serially diluted, and added in duplicate to 12-well plates following the protocol described in the MAYV Titration and Plaque Assay section.

To characterize viral cytopathic effects (CPE), 0.25% crystal violet containing 30% methanol was added to each flask for 30 minutes. To titer the harvested supernatants, plaque assays were performed on Vero cells following the same protocol as previously mentioned.

### Type I/II IFN Sensitivity Assay: Pre-treatment vs Post-treatment

To assess the role of interferon (IFN) in MAYV infection, we followed the methods of *Fros et al* (18). In brief, for IFN pre-treatment, cell lines were grown in 24-well plates to 100% confluence and IFN-β or IFN-γ was added in duplicate at 2, 20, 200, and 2000 units/ml for 6 hours. Old media containing IFN was removed and cells were washed once with DPBS w/o Ca/Mg. Medium containing MAYV-U at a MOI of 1 was then added for a 3-hr adsorption at 37°C and 5% CO_2_. Following adsorption, cells were rewashed once with DPBS, fresh media was added, and then incubated for 21 hours at 37°C and 5% CO_2_. After 21 hours, supernatants were harvested, centrifuged at 1200 xg for 10 minutes and frozen at -80°C (18). To assess viral replication in cells treated with IFN after infection, cells were infected first with MAYV-U at a MOI of 1 for a 4-hr adsorption at 37°C and 5% CO_2_. Cells were then washed with DPBS and fresh media containing IFN-β or IFN-γ was added at the aforementioned concentrations in duplicate for a 21-hr incubation. After 21 hrs, supernatants were harvested, centrifuged at 1200 xg for 10 minutes and frozen at -80°C (18). All supernatants were titered following the plaque assay protocol described earlier.

### Viral genes Sequencing

Total RNA of each isolate was extracted from frozen virus stock using QIAamp Viral RNA Mini Kit (Qiagen, Germantown, MD) following the manufacturers spin protocol. RNA concentration and quality were measured by Qubit® 2.0 (Life Technologies, Carlsbad, CA) and spectrophotometry (DeNovix® DS-11 FX). First-strand synthesis was performed with Superscript III First Strand Synthesis system (Thermo Fisher Scientific, Waltham, MA) using an oligo(dT)_20_ primer in a BIO-RAD T100™ PCR thermocycler (BioRad, Hercules, CA) using a modified protocol for viral envelope protein. Briefly, the reactions were incubated at 50°C for a total of 180 minutes, with pausing the incubation after 90 min to add an additional 2μL of SuperScript to each reaction to create full-length transcripts. The protocol otherwise followed manufacturer’s instructions. Five overlapping primer pairs for the envelope protein of the D and L genotypes were constructed using MAYV reference sequences obtained from GenBank (https://www.ncbi.nlm.nih.gov/genbank) by identifying sections of homologous base pairs with similar melting temperatures (Table S1). The cDNA template for each isolate was PCR amplified using Platinum™ Green Hot Start PCR Master Mix (2X) (Invitrogen™, Carlsbad, CA) using 3 μl of template and following the manufacturer’s instruction. The reaction was amplified using a BIO-RAD T100™ PCR thermocycler (BioRad, Hercules, CA) programmed for an initial desaturation at 94°C for 90 seconds, followed by 31 cycles using the following conditions: denaturation at 94°C for 30 seconds, annealing at 51°C, 52°C or 58°C for 30 seconds, and extension at 72°C for extension. These cycles were followed by a final extension step at 72°C for 5 min. The amplified PCR products were assessed for appropriate amplification by electrophoresis on a 1% TAE agarose gel containing 0.3μg/mL ethidium bromide. Reactions showing a single band at the appropriate molecular weight based on a 100bp ladder (Promega, Madison, WI) used for reference were purified using QIAquick PCR purification kit (Qiagen, Germantown, MD) following the manufacturer’s instructions and concentrations determined by spectrometry (Devonix® DS-11 FX, Wilmington, DE). A single site for MAYV-P (site 5) required gel extraction due to the presence of a second band on the gel. The band of interest was extracted from the gel and purified using QIAquick Gel Extraction Kit (Qiagen, Germnatown, MD) following the manufacturer’s instructions. The purified PCR reactions were prepared for Sanger sequencing and sent to Genewiz following the company’s instructions. Sequences were assessed using Geneious Prime 2019.1.3. The reverse and forward sequences for each isolate were trimmed, aligned, and inspected for congruency, missing, or uncalled base pairs and manually edited based on the chromatogram results. The five segments of envelope sequence were then assembled into a single contig with the resulting contig spanning E3, E2, 6K, and E1 for each isolate. Sequences have been deposited in GenBank, with accessions numbers: MZ962428-MZ962430.

### Genetic and Structural Analysis

In this study we analyzed E3-E2-E1 polyprotein of MAYV-U, MAYV-B and MAYV-P (Figure S1) from residue 10 of E3 to residue 423 of E1 (according to UniProt ID Q8QZ72). MAYV envelope gene sequences were used to infer the maximum likelihood phylogenetic relationship with IQ-TREE software (19) based on the best-fit model according to the Bayesian Information Criterion (BIC) (20) with ultrafast bootstrap approximation (21). Amino acid pattern along the phylogeny were identified by signature pattern analysis using VESPA (22) (https://www.hiv.lanl.gov/content/sequence/VESPA/vespa.html), while selection analyses was carried out with algorithms implemented in the HyPhy software (23) (http://www.datamonkey.org/), which estimate non-synonymous (dN) to synonymous (dS) codon substitution rate ratios (ω), where ω<1 indicates purifying/negative selection and ω>1 diversifying/positive selection. Fast Unconstrained Bayesian Approximation (FUBAR)(24) was also used for inferring pervasive selection, and the mixed effects model of evolution (MEME) (25) to identify episodic selection. Sites were considered to have experienced positive or negative selective pressure based on posterior probability (PP) > 0.90 for FUBAR, and likelihood ratio test ≤ 0.05 for MEME. The Cryo-EM structure of the mature and infective MAYV (PDB ID 7KO8, unpublished and available at https://www.rcsb.org/structure/7KO8) was used to map the mutations of interest within the 3D structure of E2 protein resolved with E1 and capsid proteins. Visualization of the atomic model, including figure and movie, is made with Chimera v1.12 (26).

## Results

### MAYV envelope is detectable in all tested target cells except for macrophages

Immunofluorescence (IF) microscopy showed that MAYV-U replicated in several tested target cells (Vero, HEK293T, fibroblasts, and SkMc) between 24 and 72 hours post infection (hpi). By 24 hpi, all samples had detectable viral envelope, the primary target of the host immune response, as indicated by the positive FITC (green) signal, which is suggestive of viral replication. As time progressed, the visible FITC signal increased, eventually peaking between 48 and 72 hpi. Within 24 hrs, infected fibroblasts had a substantial amount of detectable MAYV -E (Fig 1A), suggesting the cells were quickly infected. HEK cells also became infected within 24 hrs with some loss in cell density with increasing time, suggestive of MAYV induced cytopathic effects (Fig 1B). SkMc cells were readily infected based on detectable envelope at 24 hpi (Fig 1C). Viral envelope was not completely detectable on Vero cells at 24 hpi compared to other cell lines but was present on nearly all cells by 48 hpi, and it drastically reduced at 72 hpi due to substantial cell death as shown by the lack of DAPI positive cells (Fig 1D). While some macrophages became infected with MAYV within the first 24 hpi, infection was only minimally detectable thereafter (Fig 1E). In fibroblasts and SkMc, as time progressed, we observed condensation and fragmentation of the nuclei, known as karyorrhexis and pyknosis, respectively, which suggested that the cells were undergoing apoptosis (Figs 1A & 1C).

**Fig 1:**
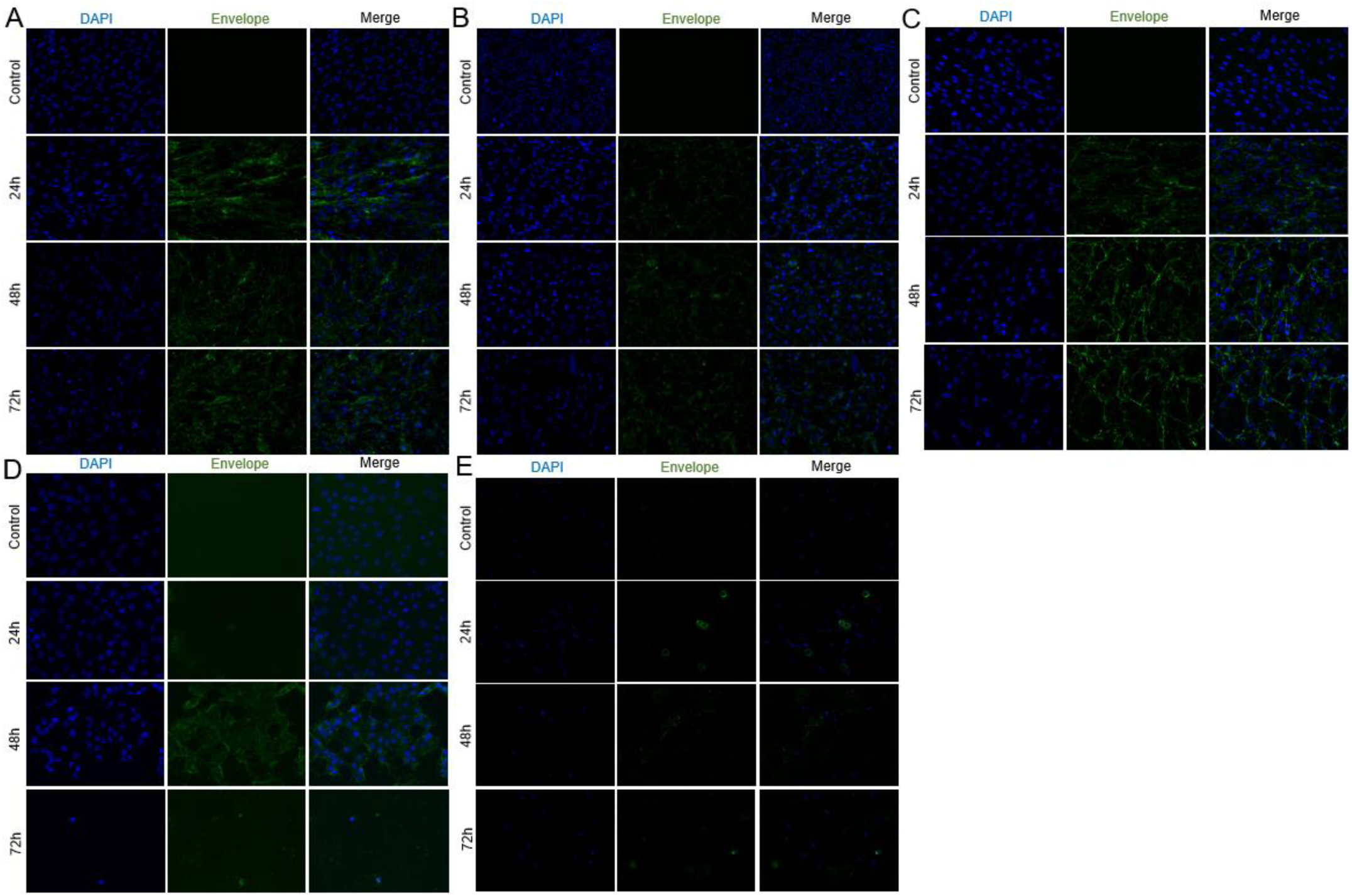
Immunofluorescence confocal microscopy of MAYV infection in target cells. Cells (**A**: Fibroblasts; **B:** HEK293T; **C:** SkMc; **D:** Vero; **E:** Macrophages) were infected with MAYV-U at an MOI=1 for 24, 48, or 72 hours. Cells were then incubated with Anti Eastern Equine Encephalitis primary antibody that is broadly specific to alphavirus envelope at 1:1000 for detection of MAYV envelope. Cells were then washed and incubated with Goat anti-Mouse IgG Alexa Fluor 488 secondary antibody at 1:1000 for primary Ab detection. The cells were lastly stained with DAPI 1:1000 and imaged at 20x using confocal microscopy. Images were Z-Staked, deconvoluted, and renormalized for processing.

### MAYV cytopathic effect varied marginally based on cell type and viral isolate

For cell lines that had significant positive IF detection (SkMc, Fibroblasts, Vero, and HEK293T), CPE was examined between viral isolates to elucidate potential differences in virulence through crystal violet staining of infected monolayers of cells. MAYV isolates ranked from greatest to least CPE as follows: MAYV-U>MAYV-B>MAYV-P. However, after qualitatively visualizing disruption of a confluent cell monolayer and changes in cell morphology, we concluded that differences were minimal. With all viral isolates, Vero cells, which have no natural innate antiviral mechanisms, showed the greatest loss in cell viability (Fig 2A). HEK293T cells, however, showed little CPE across viral isolates (Fig 2B). Fibroblasts and SkMc both demonstrated significant changes in cell morphology and monolayer disruption (Figs 2C & 2D).

**Fig 2:**
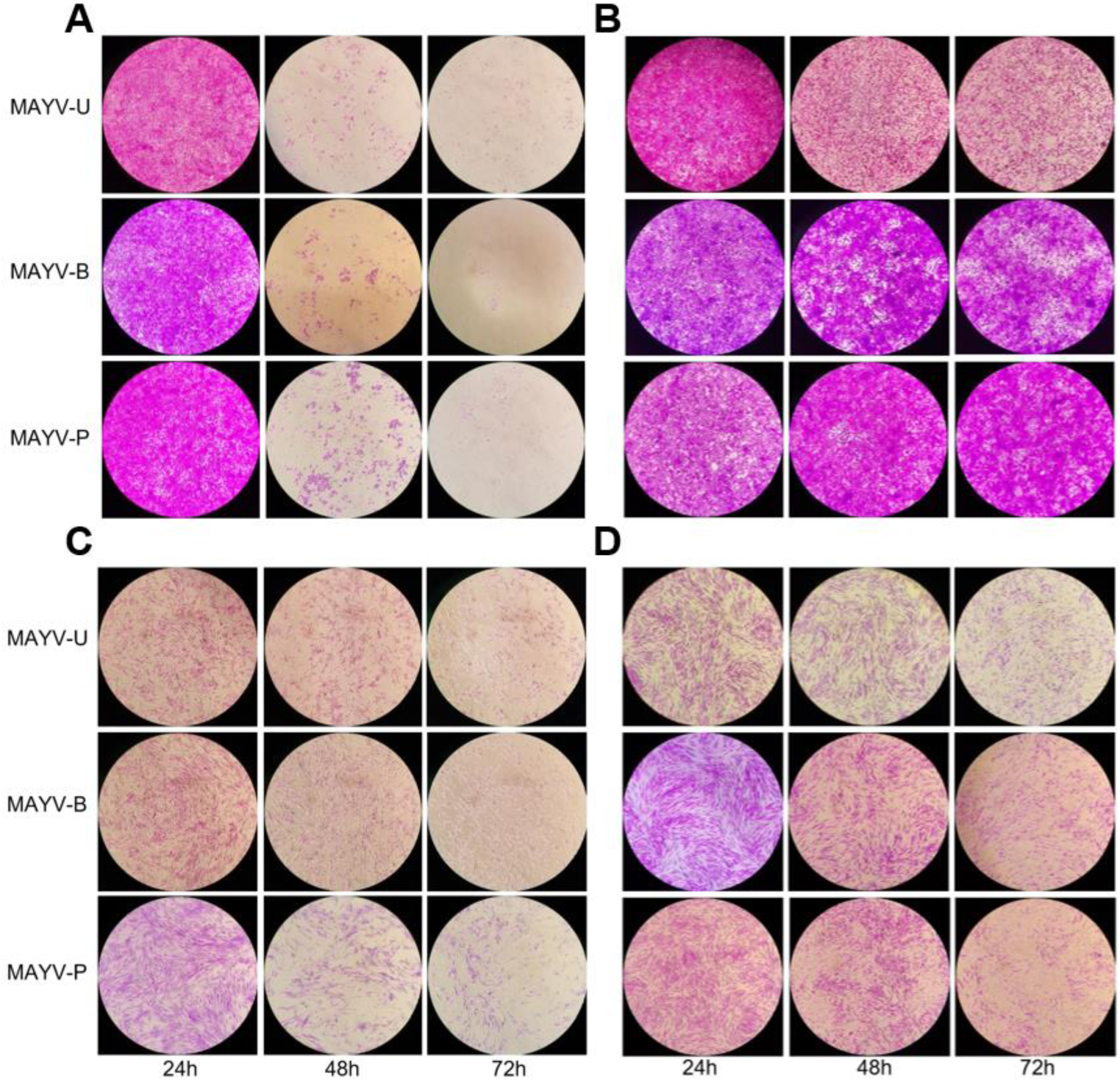
Evaluation of MAYV cytopathic effect. Cells (**A:** Vero; **B:** HEK293T; **C:** SkMc; **D:** Fibroblasts) were infected at an MOI=1 and incubated for the specified time point. Cells were then stained with crystal violet and imaged to evaluate the cytopathic effects.

### MAYV replication kinetics showed small differences between viral isolates

Comparing slopes of infection kinetics for all three strains, we found that the MAYV-U isolate possessed the greatest rate of infection, followed by MAYV-B, and MAYV-P isolates (Fig 3). These differences in kinetics do not appear to impact peak titers as each strain reaches around 10^8^ pfu/ml in all cell types tested, with peak titers occurring at approximately 30-40 hpi. Macrophage kinetics are not graphed because no increase in viral titers beyond inoculated amount was seen. When macrophages were infected with MAYV-U at an MOI=1, the viral titer at time 0 hpi was equal to 7.65×10^4^pfu/ml, which declined to 1.5×10^4^pfu/ml at 24 hpi, and further to 4.5×10^3^pfu/ml and 1.5×10^2^pfu/ml at 48 and 72 hpi, respectively. Both fibroblasts and SkMc, which are MAYV target cells, demonstrated the greatest rate of change during the first 12 hours of infection.

**Fig 3:**
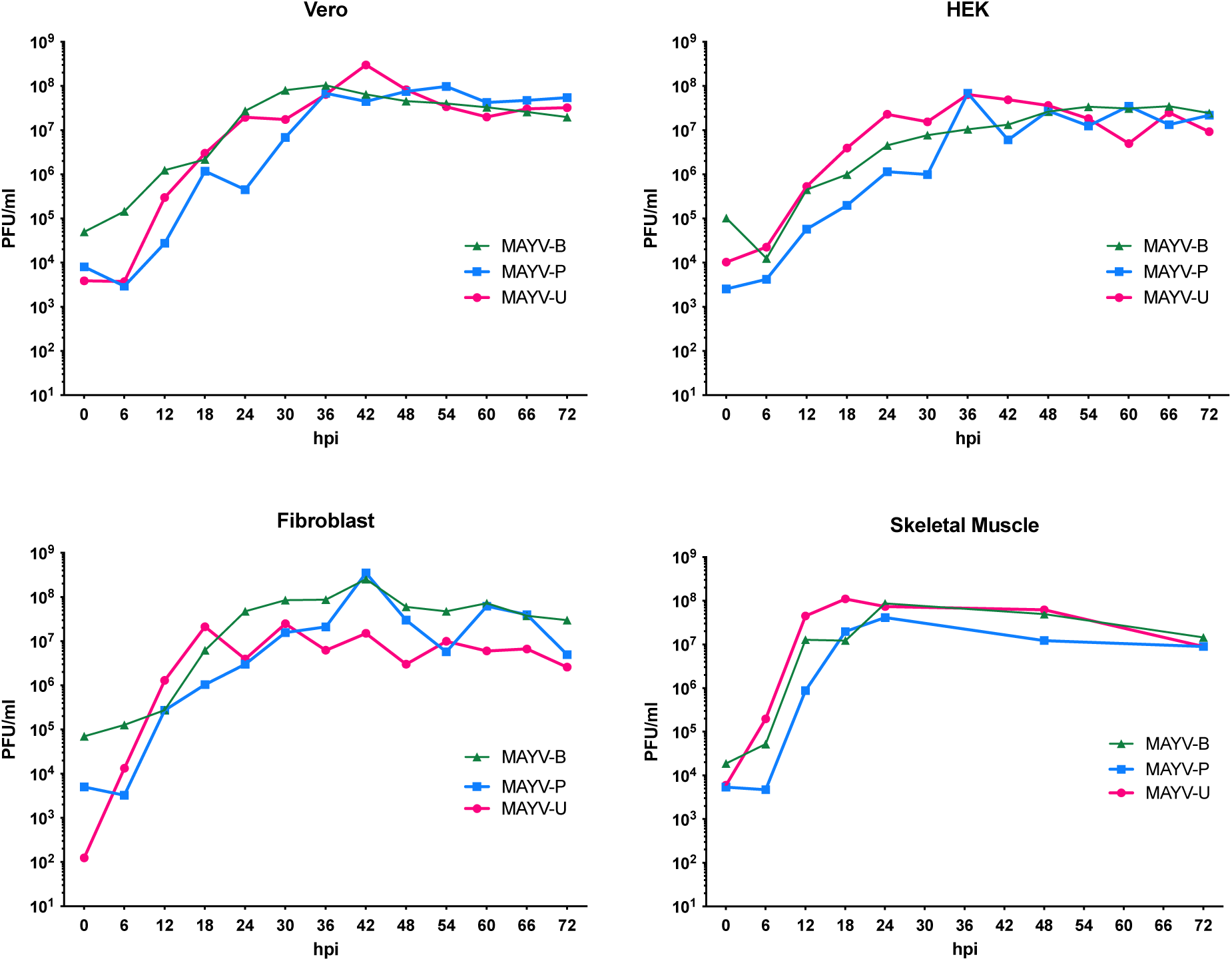
Mayaro viral replication kinetics. MAYV isolates Brazil, Uruma, Peru were inoculated at an MOI 1 on (**A:** Vero; **B:** HEK293T; **C:** Fibroblasts; **D:** SkMc) Viral titers were determined up to 72h p.i.

### Type I Interferon pre-treatment reduces viral replication

Plaque assays from both Vero cells and fibroblasts have confirmed that pretreatment with IFNβ reduces viral replication in a dose-dependent manner (Fig 4). Pretreatment with IFNγ yielded no reduction and post-treatment with either IFNβ or γ did not cause a significant reduction. HEK cells had no apparent reduction in viral titers regardless of the interferon used and timing of treatment. Treatment with IFN after infection did not cause as substantial of a reduction as pretreatment. The reduction that did occur with fibroblasts and Vero cells, though small, was dose dependent. HEK cells saw no change regardless of dose.

**Fig 4:**
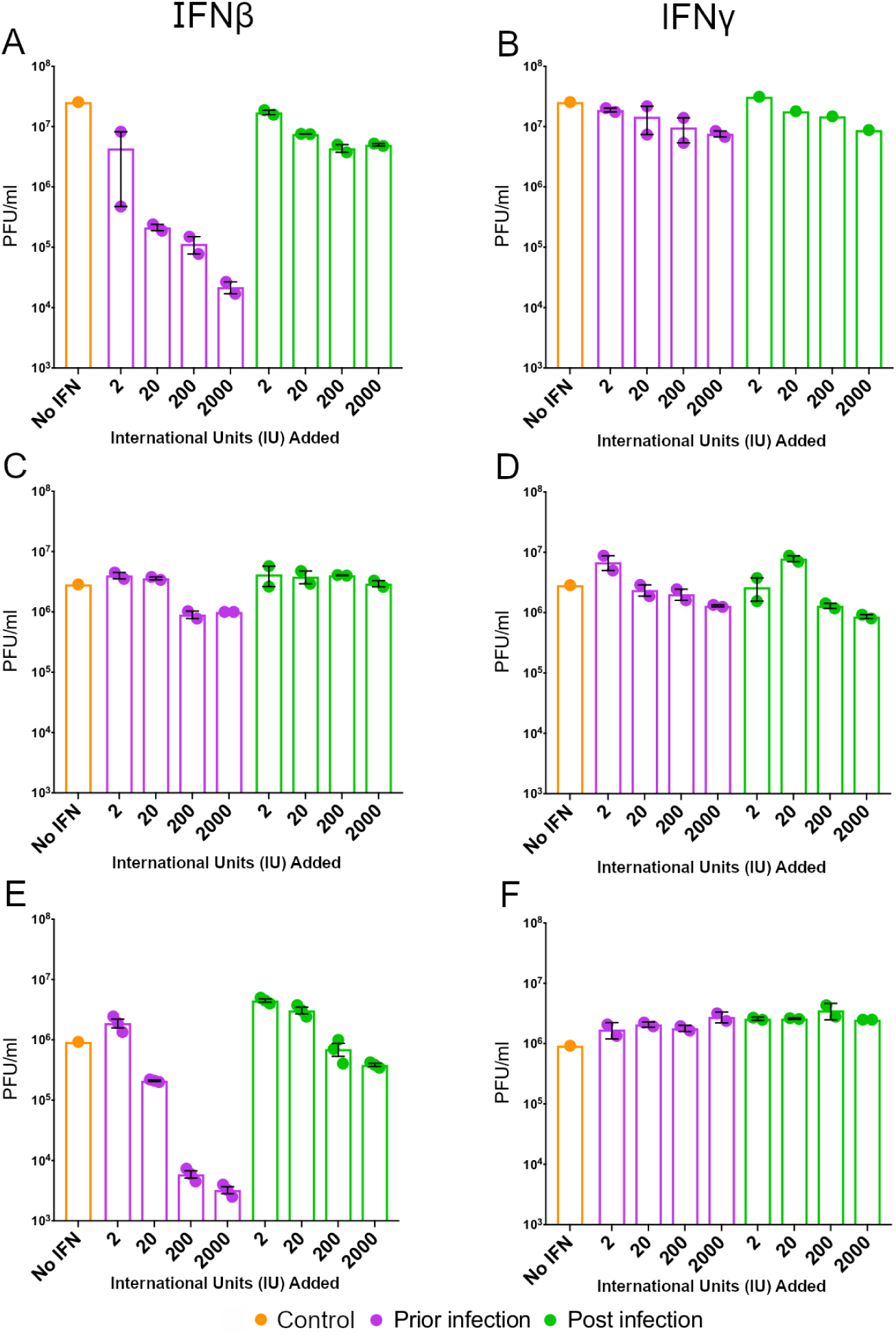
Treatment with IFNβ and IFNγ pre and post MAYV-U infection. **A-B** Fibroblasts; **C-D** HEK293T; **E-F** Vero. Cells were either infected with MAYV-U at a MOI =1 either prior to or after IFN β or γ treatment at 2, 20, 200, and 2000 IU/ml. Cell supernatants were harvested at their respective 24-hour time point and titered by plaque assay.

### Mutations in the E2 gene might influence viral fitness

Phylogeny reconstruction based on MAYV envelope genes sequenced in this study confirmed that MAYV-U (fast replication) and MAYV-P (intermediate replication) strains belong to genotype D, while MAYV-B (slow replication) strain belongs to genotype L (17) (Figure S1). Signature pattern analysis, which identifies residues unique to MAYV-U, MAYV-P and MAYV-B, showed that amino acids implicated in disulfide bonds and cleavage of the E3, E2, E1 proteins are conserved among all isolates, as well as those modified by the host at the post-transcriptional level (N-linked (GlcNAc) asparagine or S-palmitoyl cysteine modifications). However, the analysis also highlighted several key residues within the envelope region that distinguish MAYV-U from MAYV-P and MAYV-B (Figure S1). In particular, MAYV-U was the only isolate to have the following mutations in the E2 protein: the D60G mutation, responsible for switching aspartic acid for glycine in position 60, and the S205F mutation, introducing a phenylalanine instead of a serine in position 205 (Figure S1, Figure 5). Based on the experimental data, we hypothesized that these mutations may be possibly responsible for the observed *in vitro* differences in virus infection. Selection analyses, that determine whether observed frequency of mutational patterns in a specific site is expected under a neutral vs. a selection scenario (see Methods), did not find these sites to be under diversifying/positive selection. However, these mutations are of particular interest as E2 plays a role in viral attachment to the target host cell by binding the cell receptor (27). We further investigated how these mutations may affect the structure of the protein using the recently published crystal structure of E1-E2 MAYV (see Methods). The D60G mutation, which substitutes a negatively charged hydrophilic amino acid for a hydrophobic neutral amino acid, is located in the domain A; whereas S205F, that introduces a polar charged amino acid instead of an aromatic amino acid, is located in domain B of the amino-terminal region of the E2 glycoprotein (28,29). Based on the structural modeling, we conclude that the unique mutations found in MAYV-U, that map to the exposed region of the E2 glycoprotein facing cellular surface, are likely to alter the interaction of the virus with its host receptor and therefore may be related to differential fitness.

**Fig 5:**
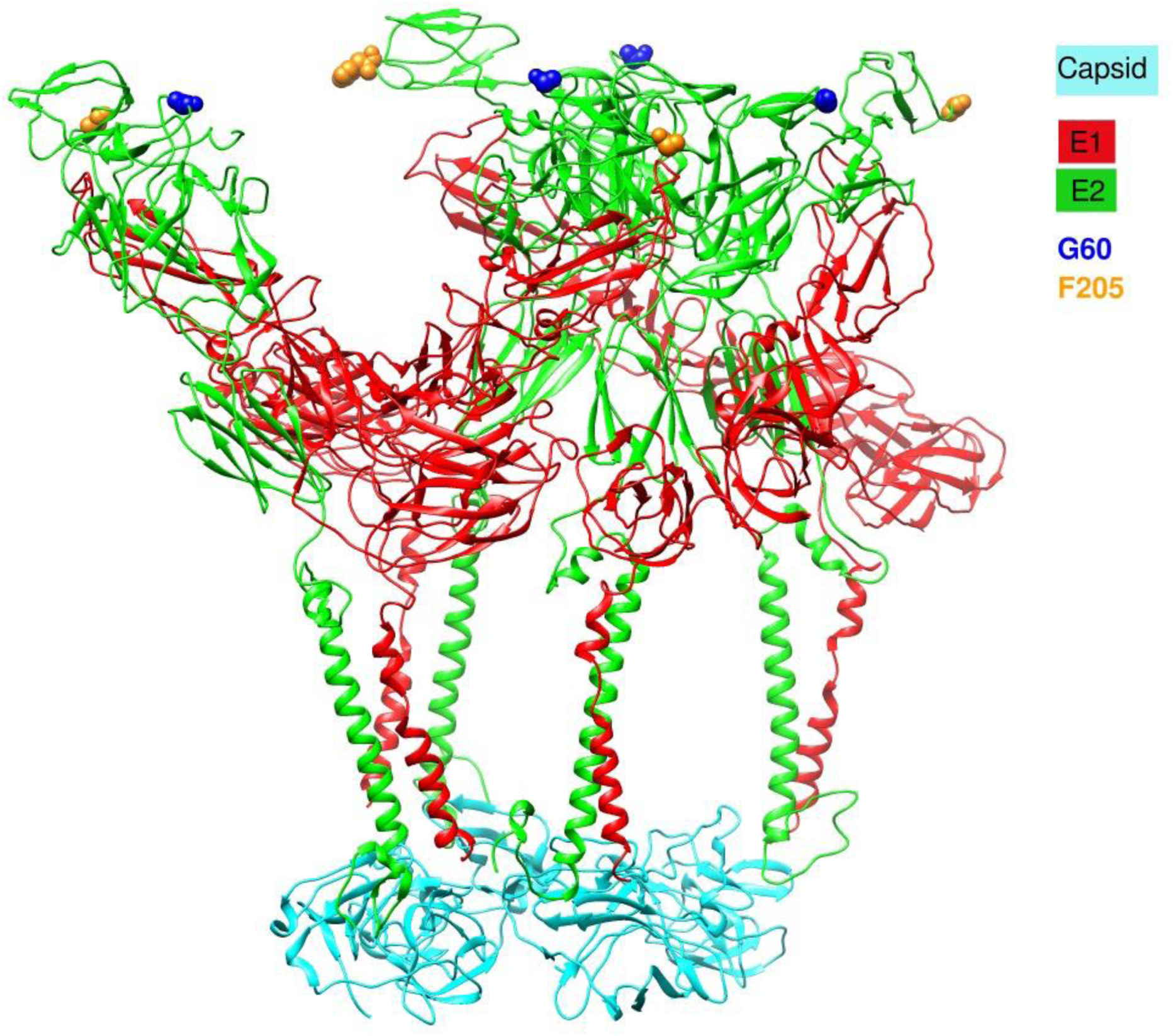
Mapping of the mutations of interest (G60 and F205) on the 3D structure of E2 protein. 3D structure of the recently available Cryo-EM structure of the mature and infective Mayaro virus (PDB ID 7KO8); G60 and F205 are represented in blue and yellow, respectively. The E1 and E2 proteins are shown in red and green, respectively, and the capsid in cyan.

## Discussion

Tropical regions in the Caribbean, Central and South America are considered high risk for MAYV emergence, yet little is currently known about MAYV pathogenesis. This is likely a result of a decreased perception of serious threat to human health. Some factors that led to this perception are MAYV’s low viremia in humans, ineffectiveness as an urban virus, limited reach as a result of sylvatic transmission, and geographic restriction to mainly Central and South American rainforests (14,30). However, CHIKV also faced similar perceptions before mutations in the envelope protein (E1-A226V) resulted in its effective replication in the widespread vector, *Aedes albopictus*, increasing its geographic range and threat to global public health (31).

We addressed these gaps in the literature by investigating the *in vitro* cytopathogenicity and viral replication kinetics of MAYV in human cell lines. Our results show that fibroblasts and SkMc are highly susceptible to MAYV infection as shown through immunofluorescent visualization of MAYV-E within 24 hpi. Infection of these cell types is important because fibroblasts are among the first cells in the body to come into contact with the virus upon inoculation from a mosquito taking a bloodmeal (32). Additionally, infection of SkMc likely contribute to the signs and symptoms that develop during infection (33). While we did not investigate cytokine secretion by infected fibroblasts and SkMc and their effects, *Bengue* et al. determined that MAYV-infected human chondrocytes have a substantially increased expression of proinflammatory cytokines IL-6 and TNF-α, both of which are mediators of arthritis (9). They genotype MAYV strain isolated in Haiti (2) in their experiments, and it remains to be understood if the results extend to MAYVD and MAYL.

In our study, Vero cells, SkMc and fibroblasts all experienced CPE, but to different degrees. Vero cells are a permissive cell line due to their inability to respond to viral infection through interferon production (34). As a result, Vero cells had rapid monolayer disruption, indicating cell death. HEK293T (human embryonic kidney epithelial cells) showed little CPE. This may be due to possible differences in interferon production. It is currently unclear whether HEK293T cells can produce Type I interferons. In addition to possible interferon production, the transformed nature of HEK239T may make them naturally more resistant to apoptotic signaling (35), therefore reducing the degree of CPE seen on infection. However, fibroblasts and SkMc cells experienced severe CPE, shown by substantial changes in cell morphology, but fewer cells detached from the flask surface compared to Vero cells. A future study would be to qualify whether these cells are still alive, characterize their cytokine expression profile, and determine viral output for extended time points.

Our viral replication kinetics experiments revealed that peak viral titers vary by cell type within a margin of one log that could be explained by the cell’s intrinsic biology and ease of viral entry/exit, etc. There is rapid replication and release of virus during the first 24 hours, which begins to slow and flatten after 24-30 hpi. Early replication was fastest for MAYV-U in all tested cell lines. While CHIKV can readily infect macrophages (36), our study showed MAYV infection is not productive in macrophages as seen by decreasing detection of viral envelope by immunofluorescence and decreasing viral titers with time. This may be explained by macrophage activation and a strong Type I interferon response that quickly dampened viral replication, rendering neighboring cells refractory to infection (37). It is also possible that MAYV can enter the cells, but it doesn’t appear to be effectively replicating within macrophages.

To address the possibility of IFN influencing MAYV replication, we performed some preliminary studies to determine whether MAYV is sensitive to the effects of interferon. The IFN sensitivity assay showed a reduction in viral titers in Vero and fibroblasts cells pretreated with IFN-β in a dose dependent manner; showing cells can respond to interferon and become more resistant to viral infection. However, this effect was not seen in IFN-β post-treated cells. This suggests that MAYV replication may be interfering with the pathway, but it is unknown how efficient or strong this response is and how it fares in an *in vivo* system with several innate immune responses acting simultaneously. For example fibroblasts are known to secrete proinflammatory and antiviral cytokines such as IL-1, IL-6 and IFN-β, which may serve as initiators of the innate immune response by activating and recruiting resident immune cells (38). It is also unknown what MAYV proteins may be responsible for the effect seen *in vitro* and these questions all call for further study. What is known, however, is that several other arboviruses such as chikungunya, dengue, Japanese encephalitis, and HHV-8 have developed interferon-evading mechanisms that aid their ability to reach high titer viremia (18,39).

Viral mutations may also explain the differences in MAYV infection between the different strains. We have shown that MAYV-U already has unique mutations in the E2 protein, D60G and S205F. The common property of the new substituted amino acids is the loss of charged residues. The domain A contains the receptor binding site, (40) whereas domain B protects the fusion loop on the domain 2 of E1 and involved in cellular attachment (28). It is possible that these mutations have the potential to enhance the host receptor binding during viral infection and facilitate viral entry into susceptible cells. A similar mutational substitution (D to G) has been reported for SARS-CoV-2, the D614G mutation (41), that alters fitness (42) and increased infectivity of the virus (43). How these mutations impact transmission remains to be studied. For example, future studies could investigate whether the signature residues distinguishing MAYV-D and MAYV-L, under diversifying selection (residue 381 in E2 protein, and 300 in E1) may increase vector susceptibility to infection and transmission and may explain the “dispersed” versus “limited” range of the genotypes. CHIKV faced limitations in invertebrate vectors and geographic range before mutations in the envelope protein (E1-A226V) enhanced its ability to infect *Aedes albopictus* (31). RNA viruses, such yellow fever virus, MAYV, and CHIKV, rapidly accumulate errors in their genomes that are transmitted to progeny virions (44). However, strong purifying selection ensures that these mutations do not weaken their fitness for their insect or mammalian hosts (16,45).

While MAYV is predominantly maintained in a sylvatic cycle with periodic incidental human infection, anthropological and environmental changes such as land use, migration, rapid urbanization, and climate change may drive MAYV toward emergence as it is forced to adapt to new susceptible hosts, mosquito vectors, and conditions (46,47). *Aedes aegypti*, a far more urbanized and anthropophilic mosquito than *Haemagogus* mosquitoes, has been shown experimentally to be a competent vector for MAYV, but low viremic titers in humans serve as a barrier to establish infection from humans into the mosquito (48,49). Increased exposure between competent vectors and MAYV may benefit the already adaptable virus, allowing transmission to become more efficient, risking the establishment of an urban cycle. This transition has occurred before in CHIKV and yellow fever, among other viruses (49,50). Evidence of this occurring in MAYV may already exists as there are three reported cases of the virus in Haiti, without the existence of *Haemagogus* mosquitoes (2,16). This may be a sign that MAYV is adapting to other vectors such as *Aedes aeqypti*. As such, we believe that our study, along with others, demonstrate the risk MAYV possesses in regard to its potential for expansion, adaptation, and suitability to several human target cells. These studies have created a strong foundation of knowledge regarding MAYV, but we believe additional epidemiological, vector ecology, and virological studies are all warranted to better understand MAYV and its risk of emergence.

## Acknowledgements

We would like to thank Aya Tal-Mason, Alexis Coican, Patricia Zielinski, and Andrew Chrystmann for their contributions and assistance in several experiments. We would also like to thank Dr. David Pascual and UTMB WRCEVA for providing the viruses used in this study.

---

**S1 Table:**
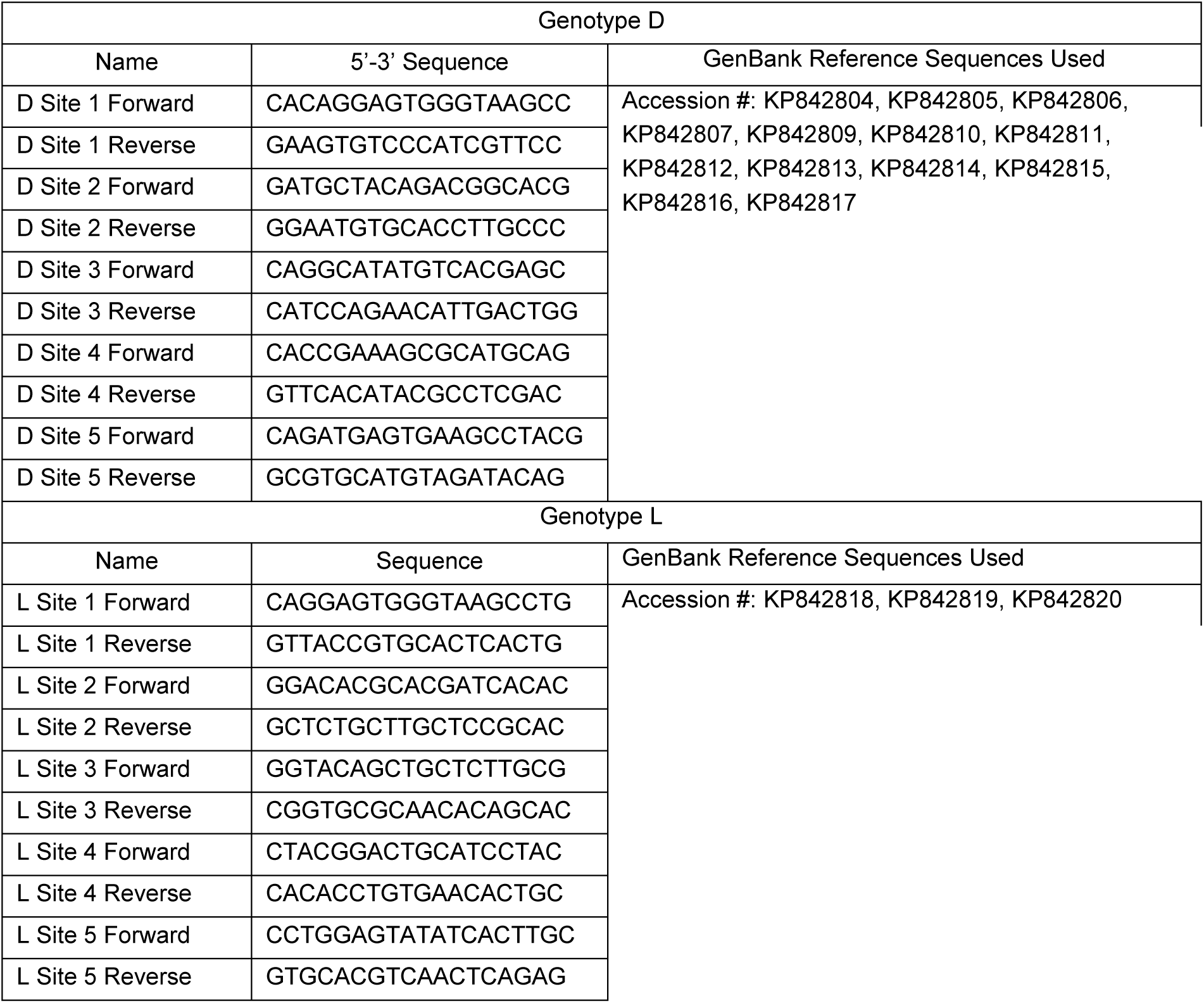
Mayaro virus primers for D and L genotype envelope protein. Five overlapping primer sets spanning the envelope protein (E3, E2, 6K, E1) for D and L, MAYV genotypes were constructed using reference sequences obtained from Genbank.

**Figure S1:**
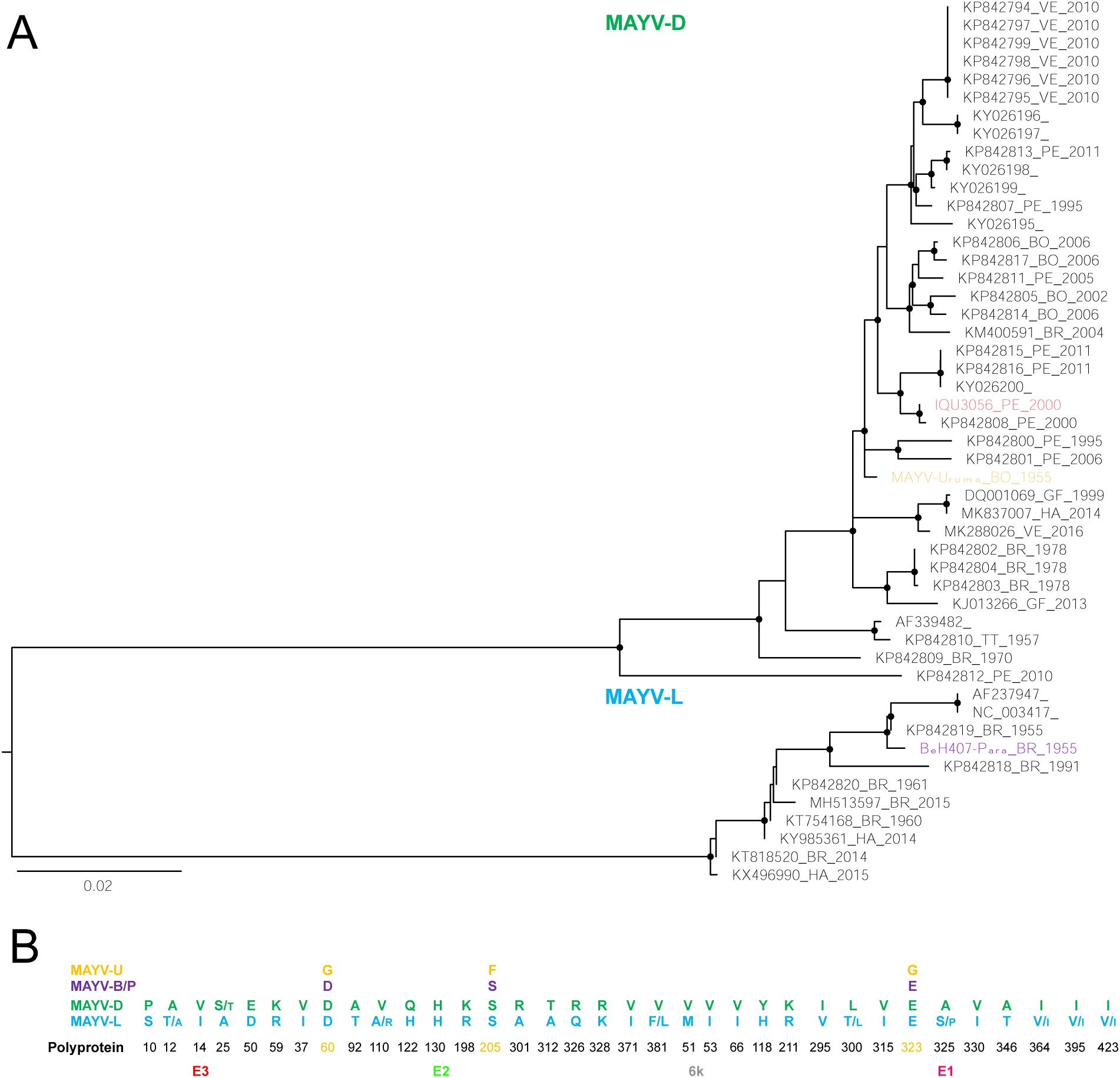
Phylogenetic inference of the structural polyprotein (E3-E2-E1) of MAYV and signature pattern analysis for MAYVL and MAYVD. Maximum likelihood phylogenetic inference of MAYV E3-E2-E1 sequences obtained from GenBank and of MAYV-U, MAYV-P or MAYV-B obtained from this study. Below the tree is given the schematic representation of the polyprotein is given and above it the blue box contains the residues that distinguish MAYVD from MAYDL as for the signature pattern analysis: the numbers correspond to the residues for each protein based on DQ487395. Above, specific residues positions of interest that are unique to MAYV-U are indicated in yellow MAYV-U, while residues found in MAYV-P and MAYV-B are in purple. In green and in blue are indicated residues found in MAYV-D and MAYD-L, respectively. Smaller letters indicate that the residue is found at lower frequency.

## Notes

### Competing Interest Statement

The authors have declared no competing interest.

